# Taxonomic replacement, but functional stability: taxonomic–functional decoupling in Amazonian streams from an Indigenous territory

**DOI:** 10.64898/2026.04.22.720173

**Authors:** Jady Vivia Almeida Silva Santos, Francieli F. Bomfim, Josinete Sampaio Monteles, Ana Beatriz Oliveira Pampolha, Juan Mateo Rivera-Perez, Paulo Geovani da Silva Gomes, Lucas Pires de Oliveira, Jair Costa Miranda-Filho, Kiarasy Kaiabi Panara, Korakoko Panara, Sewa Panara, Sakre Panara, Karapow Panara, Kwakore Panara, Sopoa Panara, Nhasykiati Panara, Pâssua Pri Panara, Pente Panara, Tepakriti Panara, Antonio Ramyllys Oliveira Costa, Lais Sarlo, Joás Silva Brito, Gabriel Martins Cruz, Raphael Ligeiro, Luciano Fogaça de Assis Montag, Karina Dias-Silva, Thaisa Sala Michelan, Leandro Juen

## Abstract

Biodiversity patterns in tropical freshwater ecosystems remain unevenly understood, particularly in high-integrity regions such as Indigenous territories. In this study, we assessed taxonomic and functional beta diversity of Ephemeroptera, Plecoptera, and Trichoptera (EPT) in Amazonian streams located within the Panará Indigenous Territory, Brazil. We evaluated the relative contributions of local environmental variables, spatial processes, and landscape context to beta-diversity patterns. We disentangled the roles of replacement and richness differences across taxonomic and functional dimensions. EPT larvae were sampled in 31 streams during the dry season. Beta diversity was quantified using Sørensen-based dissimilarity indices, and functional dissimilarity was calculated from seven ecological traits using Gower distances. Taxonomic beta diversity was dominated by genus replacement and was jointly structured by local habitat variables and spatial components, indicating the combined influence of environmental filtering and dispersal limitation. In contrast, functional beta diversity was higher than taxonomic beta diversity and was predominantly structured by richness differences, with significant effects of local environmental variables but no detectable influence of spatial processes. This pattern indicates a decoupling between taxonomic and functional dimensions, suggesting high levels of functional redundancy among EPT genera across streams. Our findings demonstrate that Amazonian streams within Indigenous territories provide key systems for understanding community assembly processes under low levels of direct anthropogenic disturbance. By revealing contrasting mechanisms underlying taxonomic and functional beta diversity, this study underscores the importance of integrating multiple facets of biodiversity and reinforces the role of Indigenous territories as strategic landscapes for safeguarding Amazonian freshwater biodiversity.

## Introduction

Despite its central role in regulating biogeochemical cycles, hydrological dynamics, and climate stability across regional and global scales, Amazonian biodiversity remains unevenly documented, with substantial gaps in remote regions where logistical constraints limit systematic surveys (Malhi et al., 2008; Intergovernmental Panel on Climate Change, 2021; Carvalho et al., 2023). These knowledge shortfalls are particularly critical in aquatic ecosystems, which are highly dynamic and structured by strong interactions between terrestrial and aquatic compartments, a defining feature of tropical floodplain landscapes (Junk et al., 1989; Ward et al., 1989; Tockner et al., 2010).

In this context, Amazonian aquatic systems play a crucial role in maintaining biodiversity and ecosystem processes, yet are increasingly exposed to pressures from climate change and land-use change (Malhi et al., 2008). Therefore, Indigenous territories represent important areas for biodiversity conservation, often maintaining lower deforestation rates and higher ecological integrity than unprotected regions (Façanha et al., 2026; Carvalho et al., 2023; Osborne et al., 2024).

These territories can be understood as socioecological refugia, where traditional management practices help maintain environmental integrity and biological diversity (Azevedo-Santos et al., 2019; Silva et al., 2025; Guerrero-Moreno et al., 2025). However, despite their high ecological integrity, these territories are not exempt from anthropogenic pressures, particularly those originating from surrounding areas, which may compromise their ecological conditions over time (Associação IAKIÔ; ISA, 2018). Even so, ecological studies conducted in Indigenous territories remain scarce, and freshwater ecosystems in these territories are underrepresented in the literature, particularly in studies that integrate multiple dimensions of biodiversity (Ceron & Silva, 2023; Carvalho et al., 2023; Santos et al., 2024). This limitation constrains our ability to evaluate how ecological processes operate under conditions of low anthropogenic disturbance, where theoretical expectations may be more clearly expressed.

Addressing these knowledge gaps, participatory monitoring approaches have gained prominence as promising strategies for generating ecological data in remote regions of the Amazon (Pereira et al., 2022). These approaches involve collaboration between researchers and local communities in the collection and interpretation of environmental data, expanding spatial and temporal coverage while incorporating local ecological knowledge into the understanding of environmental dynamics. In Indigenous territories, such approaches are particularly relevant, as they not only advance scientific knowledge but also strengthen territorial governance and community autonomy in managing natural resources (Chiaravalloti et al., 2018).

Research conducted in Indigenous territories offers a unique opportunity to examine community assembly patterns that approximate conditions close to ecological integrity, thereby advancing our understanding of the mechanisms structuring aquatic communities (Azevedo-Santos et al., 2019). Furthermore, the incorporation of participatory research strategies involving Indigenous communities represents both a methodological and ethical advancement, promoting knowledge co-production, valuing local knowledge systems, and strengthening conservation strategies aligned with socioenvironmental justice (Mateo-Vega et al., 2017; Osborne et al., 2024).

Building on this perspective, understanding how aquatic communities are structured across landscapes requires considering the interplay between local environmental conditions, ecological integrity, and spatial processes operating at multiple scales (Heino et al., 2015; Barrilli et al., 2024; Castro et al., 2020). Environmental variables can constrain species occurrence by imposing physiological and ecological limits, while dispersal capacity and spatial barriers influence access to suitable habitats (Mozzaquattro et al., 2020). When these conditions do not meet species requirements, species may decline in abundance, shift to more favorable habitats, or undergo local extinction (Monteles et al., 2021; Silva Santos et al., 2023). These processes are widely discussed within the metacommunity framework, which integrates environmental filtering, dispersal, and stochasticity in shaping biological communities (Hubbell, 2001; Leibold et al., 2004), and are reflected in variation in community composition among sites, known as beta diversity (Whittaker, 1960). In high-integrity systems such as Indigenous territories, reduced anthropogenic filtering may enhance the detectability of these mechanisms by minimizing confoundings disturbances and allowing ecological processes to operate more clearly, providing a valuable setting to test metacommunity predictions.

Beta diversity can be partitioned into two complementary components: replacement and richness differences, which represent distinct forms of variation among communities (Podani et al., 2011). The replacement component reflects species exchange along environmental or spatial gradients, whereas the richness difference component reflects gains or losses of species among sites. In aquatic ecosystems, the relative contribution of these components provides important insights into the roles of environmental heterogeneity, spatial connectivity, and dispersal limitation (Soininen et al., 2018).

To more comprehensively understand how species are distributed along environmental gradients, functional diversity has emerged as a complementary approach to taxonomic analyses. This approach is based on the characterization of species’ morphological, physiological, and behavioral traits, which are directly linked to their ecological functions (Thorp & Covich et al., 2010). These traits reflect species performance and their contribution to ecosystem processes, allowing the assessment of community resistance and resilience to environmental disturbances and global change (Santos et al., 2024; Mammola et al., 2021). Thus, integrating taxonomic and functional beta diversity provides a more robust and mechanistic analytical framework for understanding the processes shaping metacommunity organization (Laureto et al., 2015; Hepp et al., 2023).

Recent studies show that local environmental conditions directly influence functional diversity by acting as filters that select specific combinations of traits associated with physiological tolerance, resource use, and life-history strategies (Southwood, 1977; Heino et al., 2013; Cadotte et al., 2011). In contrast, taxonomic beta diversity tends to respond more strongly to spatial heterogeneity and landscape configuration, reflecting both species turnover along environmental gradients and dispersal limitations (Soininen et al., 2007; Gronroos et al., 2013; Heino et al., 2015). In some contexts, functional diversity may exhibit more conservative patterns, with high similarity among sites despite variation in taxonomic composition, indicating a potential decoupling between these biodiversity dimensions (Mouchet et al., 2010; Villéger et al., 2013). Under such conditions, functional redundancy may buffer trait composition against species turnover, leading to a greater contribution of richness differences to functional beta diversity relative to replacement.

Among aquatic organisms, insects from the orders Ephemeroptera, Plecoptera, and Trichoptera (EPT) are widely recognized for their sensitivity to habitat alterations, life cycles involving both aquatic and terrestrial phases, and varying dispersal abilities, making them a model group in aquatic metacommunity studies (Brasil et al., 2020; Callisto et al., 2023; Faria et al., 2024; Maués-Silva et al., 2024). These organisms exhibit high taxonomic and functional diversity, strong dependence on specific environmental conditions, and occupy different trophic groups, making them particularly suitable for investigating ecological patterns in Amazonian streams (Rosenberg & Resh, 1993; Faria et al., 2021; Lima et al., 2022).

Understanding the structuring of aquatic communities in protected areas is essential for elucidating ecological processes and informing conservation strategies. Despite advances in understanding beta diversity in tropical lotic ecosystems, studies integrating taxonomic and functional approaches in high-integrity environments such as Amazonian Indigenous territories remain scarce (Azevedo-Santos et al., 2019; Brasil et al., 2021). These areas provide a unique opportunity to investigate community assembly patterns under low anthropogenic pressure.

In this context, this study aims to assess taxonomic and functional beta diversity (total, replacement, and richness differences) of EPT communities in Amazonian streams within the Panará Territory. Specifically, we evaluate the relative contributions of local environmental, landscape, and spatial variables to the structuring of these patterns. We test two central hypotheses: (I) taxonomic beta diversity is jointly structured by environmental and spatial components, whereas functional beta diversity is predominantly driven by local environmental filtering; and (II) taxonomic beta diversity is dominated by the replacement component, while functional beta diversity is primarily associated with richness differences, reflecting a potential decoupling between these biodiversity dimensions. By integrating these approaches within a high-integrity socioecological context, this study provides empirical insights into metacommunity organization and highlights the role of Indigenous territories in maintaining the functional structure of Amazonian aquatic ecosystems.

## Material and Methods

### Study area

The study was conducted in 31 streams within the Panará Indigenous Territory, spanning the municipalities of Altamira (Pará), Guarantã do Norte (Mato Grosso), and Matupá (Mato Grosso) (Figure 1). The Iriri River predominantly drains the region and is the main tributary of the Xingu Basin. Vegetation is composed of Open Ombrophilous Forest, Dense Ombrophilous Forest, and transition zones between Savanna and Seasonal Forest. The regional climate is classified as tropical humid (Aw) according to the Köppen classification (Köppen, 1936), with mean annual precipitation ranging from 1,500 to 2,000 mm (Associação IAKIÔ; ISA, 2018).

**Figure 1.**
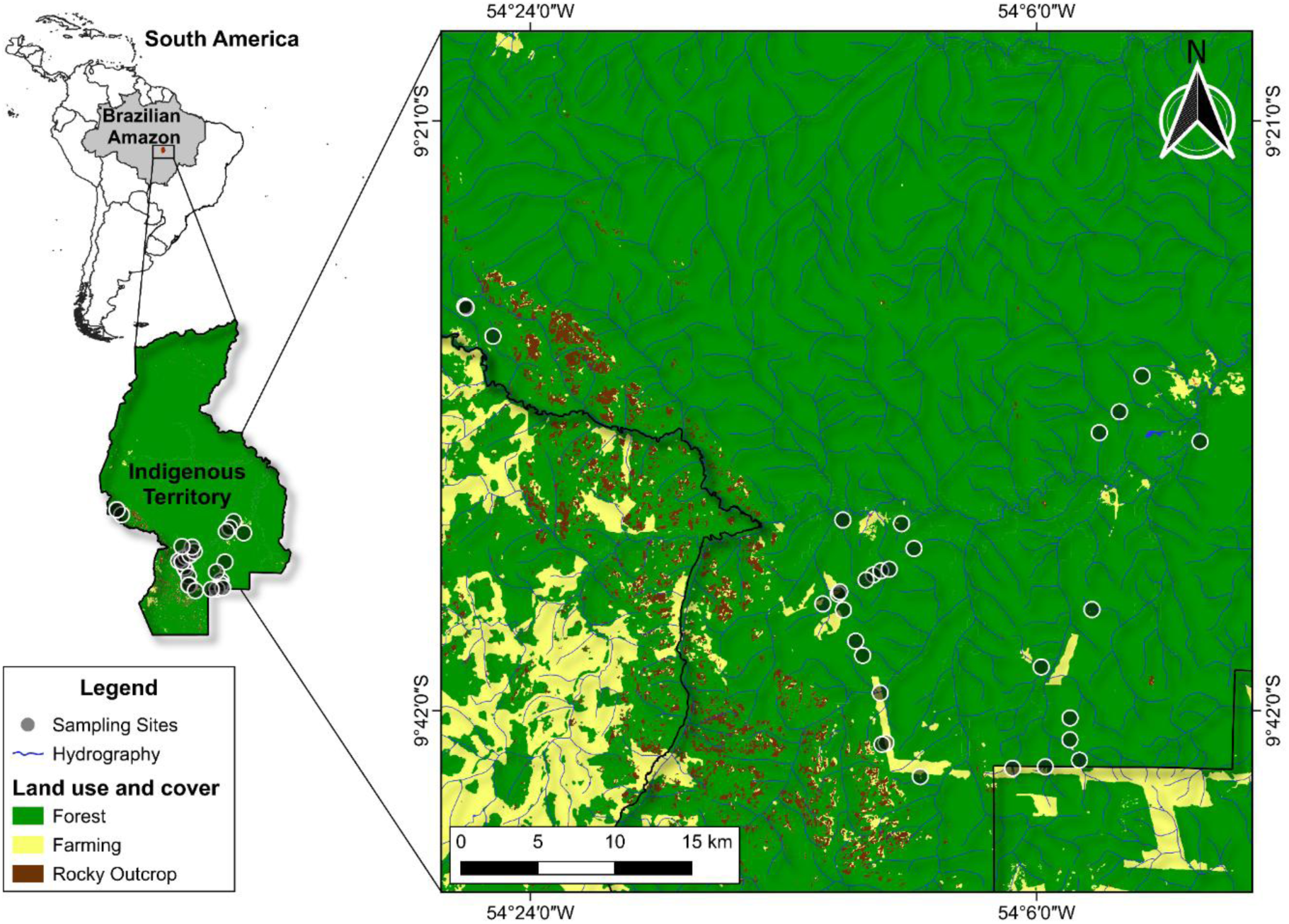
Map showing the sampling sites in the Indigenous territory, covering the municipalities of Guarantã do Norte (Mato Grosso State) and Altamira (Pará State) in Brazil.

The Panará people have a history marked by territorial displacement, intensified from the 1970s onward with the expansion of colonization fronts in the Amazon and the construction of the BR-163 highway (Cuiabá–Santarém), which crossed their traditional territory. This process led to forced contact, epidemics, and the relocation of survivors to the Xingu Indigenous Park in 1975. After approximately two decades, Panará leaders initiated a return to their ancestral lands, culminating in the reoccupation of part of their traditional territory in the late 1990s and the subsequent official demarcation of the Panará Indigenous Territory, which is currently undergoing socioenvironmental reconstruction (Moutinho et al., 2025).

Despite maintaining extensive remnants of native forest and high ecological integrity, the territory still bears marks of past land use (Associação IAKIÔ; ISA, 2018). Residual pastures persist in some internal areas, while the territory’s edges, particularly in the southern portion where proximity to agricultural production areas is greater, are more strongly influenced by surrounding farming activities, resulting in a mosaic of land uses and land-cover types (Pignati et al., 2017). In this context, streams in the region are embedded within landscapes exhibiting varying degrees of anthropogenic influence, generating environmental gradients that are relevant for structuring aquatic communities (Rocha et al., 2024).

### Ethical approval of the research

The capture, collection, and transport of biological material were authorized by the Instituto Brasileiro do Meio Ambiente e dos Recursos Naturais Renováveis (IBAMA; SISBIO permit No. 4681-1) and approved by the Animal Use Ethics Committee of the Federal University of Pará (CEUA No. 8293020418).

### Authorization for access to indigenous territory

Authorization to access Indigenous territories was granted by the Fundação Nacional dos Povos Indígenas (FUNAI) under administrative process No. 08620.012263/2023-71.

### Streams microhabitat variables

To assess stream environmental conditions, a 150-m reach was established at each sampling site, subdivided into 11 equidistant transects labeled from “A” (downstream) to “K” (upstream), defining 10 longitudinal sections between transects. These procedures followed the United States Environmental Protection Agency protocol (US-EPA; Peck et al., 2006), adapted for tropical regions (Callisto et al., 2014; Juen et al., 2025).

At each transect, the following physical habitat variables were measured: channel width (m), using a measuring tape; depth (cm), measured with a graduated ruler at representative points within each section; and canopy cover (%), estimated using a spherical densiometer or a standardized visual assessment. Substrate composition was determined through visual estimation of the relative proportion of sand, fine sediment, and organic matter, considering the wetted area of each section. Habitat integrity was assessed using the Habitat Integrity Index (HII; Nessimian et al., 2008), which integrates riparian vegetation attributes, channel structure, and substrate characteristics.

Additionally, limnological variables, including dissolved oxygen (mg L⁻¹), water temperature (°C), pH, electrical conductivity (µS cm⁻¹), and turbidity (NTU), were recorded at three points along each reach (beginning, middle, and end; transects A, F, and K) using a multiparameter probe (Horiba U-50; Asko, São Leopoldo, RS, Brazil). Limnological variables were measured prior to biological sampling (EPT immatures) and physical habitat assessment to avoid any alteration in water chemistry resulting from sampling disturbance (Maués-Silva et al., 2024).

### Spatial variables

Spatial variables were obtained using distance-based Moran’s Eigenvector Maps (dbMEM; Legendre et al., 2012), based on the Euclidean distance matrix among the geographic coordinates of sampled streams. The resulting eigenvectors were used as explanatory spatial variables in canonical ordination models. The dbMEM approach allows the detection of spatial structures across multiple scales. Analyses were performed using the dbmem function in the “adespatial” package (Dray et al., 2025) within RStudio (version 4.0.4).

### Land use and land cover variables

To obtain land use and land cover variables, we first delineated the boundaries of each stream micro catchment from the geographic coordinates of the downstream point (section “A”) using digital elevation models (INPE Topodata) in QGIS 3.34, through flow accumulation and drainage algorithms. The delineated micro catchments were visually inspected to ensure consistency with the hydrographic network.

Land use and land cover data were obtained from MapBiomas Collection 8 (2024) and verified using Landsat 5 TM imagery (30-m spatial resolution; NASA/USGS) to ensure classification accuracy within each micro catchment. For this study, land use was categorized into two dominant classes: (i) forest, including native perennial and alluvial forest formations; and (ii) agriculture, primarily associated with maize cultivation. The proportion of each class within the micro catchments was calculated and used as a landscape-scale predictor in subsequent analyses.

### Biological sampling

Each longitudinal section was subdivided into three 5-m segments; however, only the first two segments were sampled, as sampling the third segment could disturb the substrate of the subsequent section (Faria et al., 2017). Sampling was conducted during the dry season, when substrate stability is higher, reducing the likelihood of larval drift (Faria et al., 2024; Schulting et al., 2019).

Within each segment, two substrate samples were collected using a hand net (rapiché) with an 18-cm diameter and 250-µm mesh size. Each sampled segment yielded one subsample, for a total of 20 subsamples per stream (Juen et al., 2016; Faria et al., 2017; Shimano et al., 2018). Subsamples from each stream were subsequently pooled into a single composite sample, representing the EPT diversity of each sampling unit.

Collected EPT specimens were sorted and preserved in 85% ethanol. In the laboratory, individuals were identified to genus level using specialized identification keys (Dominguez et al., 2006; Hamada et al., 2014). After identification, specimens were deposited in the aquatic insect collection of the Laboratory of Ecology and Conservation (LABECO) at the Federal University of Pará.

### Functional traits

The selection of functional traits for EPT genera was based on Santos et al. (2024), which compiled a dataset of genera with distribution records in the Amazon. In total, seven traits were selected to represent EPT functional diversity: (i) body size (body width, cm), related to energetic efficiency, predation vulnerability, and dispersal capacity; (ii) trophic functional group (shredder, collector-gatherer, collector-filterer, scraper, and predator), associated with organic matter processing and trophic position; (iii) locomotion type (swimmer, sprawler, burrower, and clinger), reflecting microhabitat use and substrate interaction; (iv) body shape (flattened, cylindrical, or streamlined), related to resistance to hydraulic stress; (v) body flexibility, associated with the ability to occupy microhabitats under different flow regimes; (vi) respiration type (branchial or tegumentary), indicative of tolerance to variation in dissolved oxygen availability; and (vii) life cycle/voltinism (univoltine or multivoltine), related to population resilience to hydrological disturbances, as well as resistance strategies such as refuge use, substrate attachment, or burrowing behavior during high-flow events, when such information was available.

These traits were selected for their ability to represent ecological mechanisms related to environmental filtering, habitat use, hydraulic resistance, tolerance to limnological variation, and responses to disturbance, dimensions widely recognized as structuring aquatic macroinvertebrate communities (Poff et al., 2006; Dolédec & Bonada, 2013; Heino et al., 2015).

### Participatory sampling with Panará researchers

This study was developed through a participatory approach, involving collaboration with Panará Indigenous researchers, who participated in both training and field data collection. The involvement of local communities in biodiversity monitoring initiatives has been widely recognized as an important strategy for conservation and sustainable resource use (Corrigan et al., 2018). In Brazil, initiatives such as the ICMBio Aquatic Monitoring Program (Normative Instruction No. 3/2017) and research networks such as the National Institute of Science and Technology for Amazon Biodiversity Synthesis (INCT-SimBiAm) have promoted participatory approaches in environmental monitoring (Resende et al., 2025).

Before field sampling, we conducted training sessions with Panará researchers, including illustrated materials (slides and photographs) on aquatic insects and their role as bioindicators. Training also included practical activities in streams, demonstrating the sampling procedures used in this study, such as biological collection and environmental data recording.

Following training, biotic and abiotic sampling was conducted collaboratively with Panará researchers at the designated sampling sites (Figure 2). During fieldwork, participants helped collect aquatic macroinvertebrates and record environmental variables. This approach enabled the generation of ecological data while strengthening processes of knowledge co-production and local engagement in environmental monitoring.

**Figure 2.**
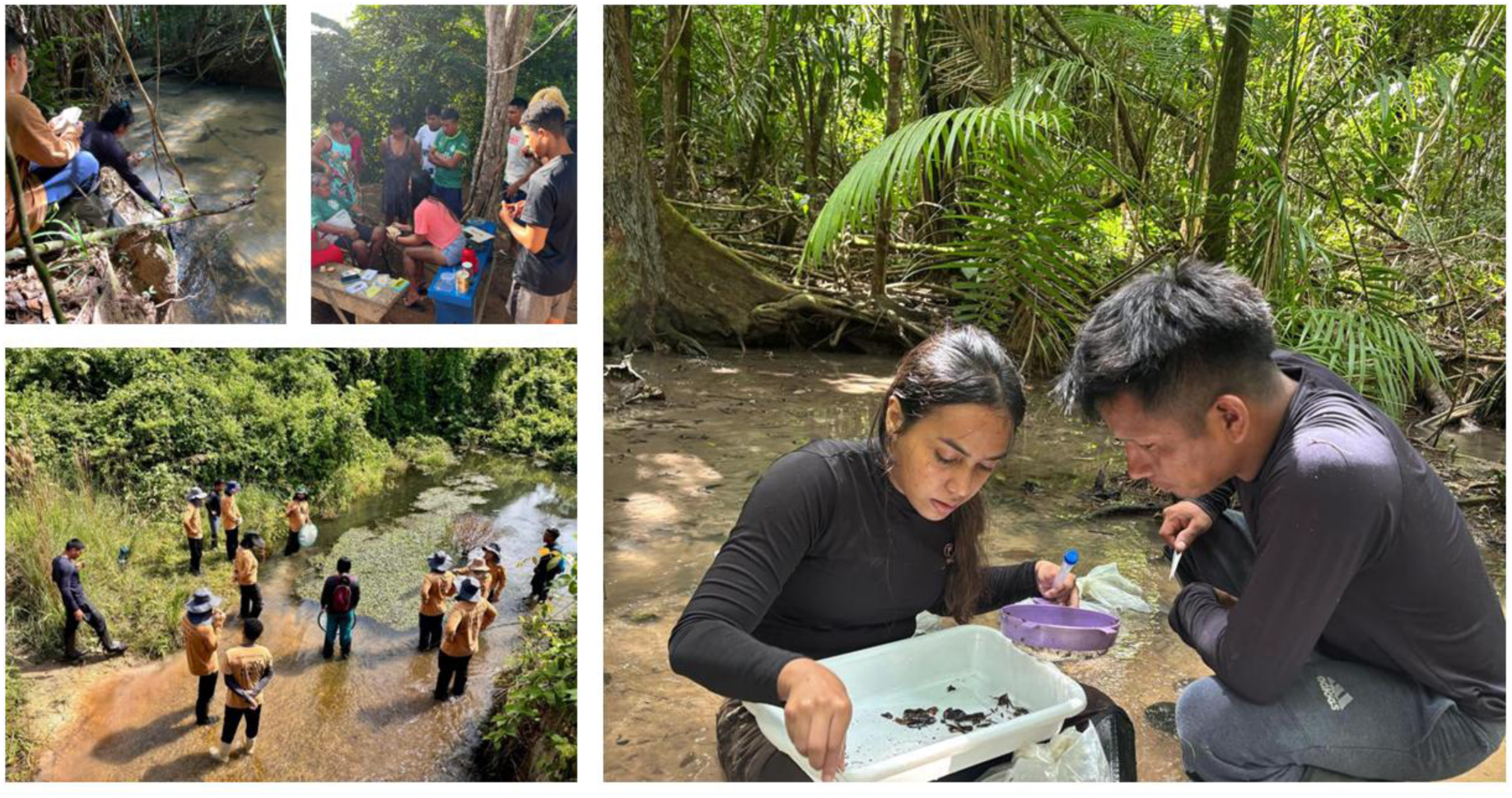
Collaborative monitoring of aquatic biodiversity with Panará Indigenous researchers.

### Data analysis

Each stream was considered a sampling unit, for a total of 31 sampling units. Taxonomic and functional beta diversity were calculated using Sørensen dissimilarity, based on EPT abundance data previously standardized using the Hellinger transformation (Legendre & Gallagher, 2001). Total beta diversity (β_total) was then partitioned into replacement (β_repl) and richness difference (β_rich) components, following Podani and Schmera (2011) and Carvalho et al. (2012), using the beta function and the packages “FD” (Laliberté et al., 2014), “BAT” (Cardoso et al., 2020), and “stats” (Bolar, 2019).

Before functional beta diversity analysis, the functional trait matrix was transformed into a Gower distance matrix (function gowdis) and subsequently converted into a hierarchical clustering structure (functional dendrogram) using the hclust function from the “BAT” package. Gower distance is a robust dissimilarity measure that efficiently handles mixed data types, allowing the integration of continuous and categorical variables. The resulting functional dendrogram, combined with the standardized abundance matrix, was used to calculate functional beta diversity.

To assess the relative contributions of environmental (microhabitat and limnological), spatial, and landscape variables in explaining the different facets of beta diversity and their components (Borcard & Legendre, 2002; Peres-Neto et al., 2006), we performed variation partitioning based on distance-based redundancy analysis (db-RDA). This analysis was used because the response variables were dissimilarity matrices derived from beta diversity components (total, replacement, and richness difference). Analyses were conducted using the capscale and varpart functions in the “vegan” package (Oksanen et al., 2019). The significance of predictors was tested using ANOVA (function anova).

Before db-RDA, multicollinearity among variables was assessed using the variance inflation factor (VIF), and variables with VIF < 10 were retained (using the function vif.cca). Predictors were selected using forward selection (function forward.sel; Blanchet et al., 2008). The spatial component was represented by Moran’s eigenvector maps (MEMs); the landscape component by the percentage of natural forest and agricultural cover; and local environmental variables by limnological and microhabitat descriptors. All analyses were conducted in R (version 4.4.0).

## Results

### EPT community composition

A total of 3,417 EPT specimens were recorded. Of these, 773 individuals belonged to Ephemeroptera, distributed across 20 genera. Plecoptera was represented by 30 individuals, all belonging to a single genus. Trichoptera comprised 2,614 individuals across 14 genera. The most abundant genera within each order were *Leptonema* (n = 459), *Miroculis* (n = 286), and *Anacroneuria* (n = 30).

### EPT beta diversity decomposition

Beta diversity decomposition revealed contrasting patterns between taxonomic and functional facets. Total taxonomic beta diversity showed a moderate magnitude (β_total = 0.539) and was predominantly driven by species turnover (β_repl = 0.488), while the richness difference component contributed minimally (β_rich = 0.051). In contrast, total functional beta diversity was high (F-β_total = 0.648) and strongly driven by functional richness differences (F-β_rich = 0.467), with a smaller contribution from functional replacement (F-β_repl = 0.180) (Figure 3).

**Figure 3.**
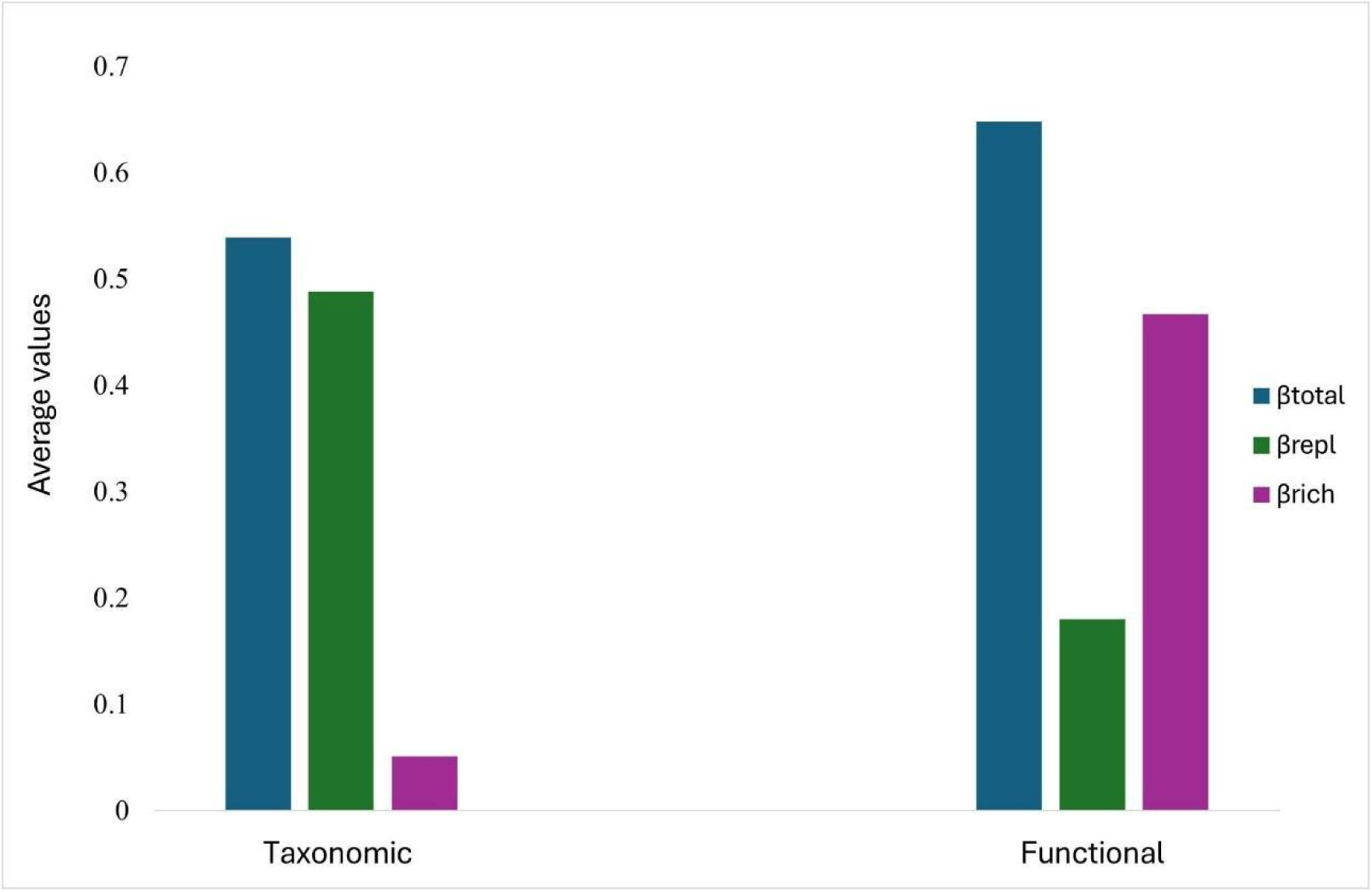
Taxonomic and Functional beta diversity values (average). β_total = total beta; β_repl = replacement; β_rich = richness difference.

### Drivers of taxonomic β-diversity of EPT communities

Both local environmental variables and spatial components significantly explained the total taxonomic beta diversity of EPT. Habitat-related variables (H) *explained* 13% of the observed variation (F = 1.948; p < 0.001), with organic matter (OM), fine substrate fraction (x.fine), and wetted width (WW) identified as the main predictors. Limnological variables (L) accounted for 7% of the variation, with water temperature (WT) as a significant explanatory variable (F = 2.270; p = 0.018). The spatial component, represented by MEM1 and MEM4, explained 14% of the total variation (F = 2.246; p < 0.001).

The taxonomic replacement (β_repl) was also significantly influenced by spatial factors (MEM1 and MEM4), which explained 16% of the variation (F = 1.604; p = 0.041), and by habitat-related variables (H), particularly organic matter (OM) and fine substrate fraction (x.fine), accounting for 11% of the observed variation (F = 1.831; p = 0.012). In contrast, the richness difference component (β_rich) was explained exclusively by the limnological variable water temperature (WT), which accounted for 37% of the variation (F = 2.284; p = 0.042) (Figure 4).

**Figure 4.**
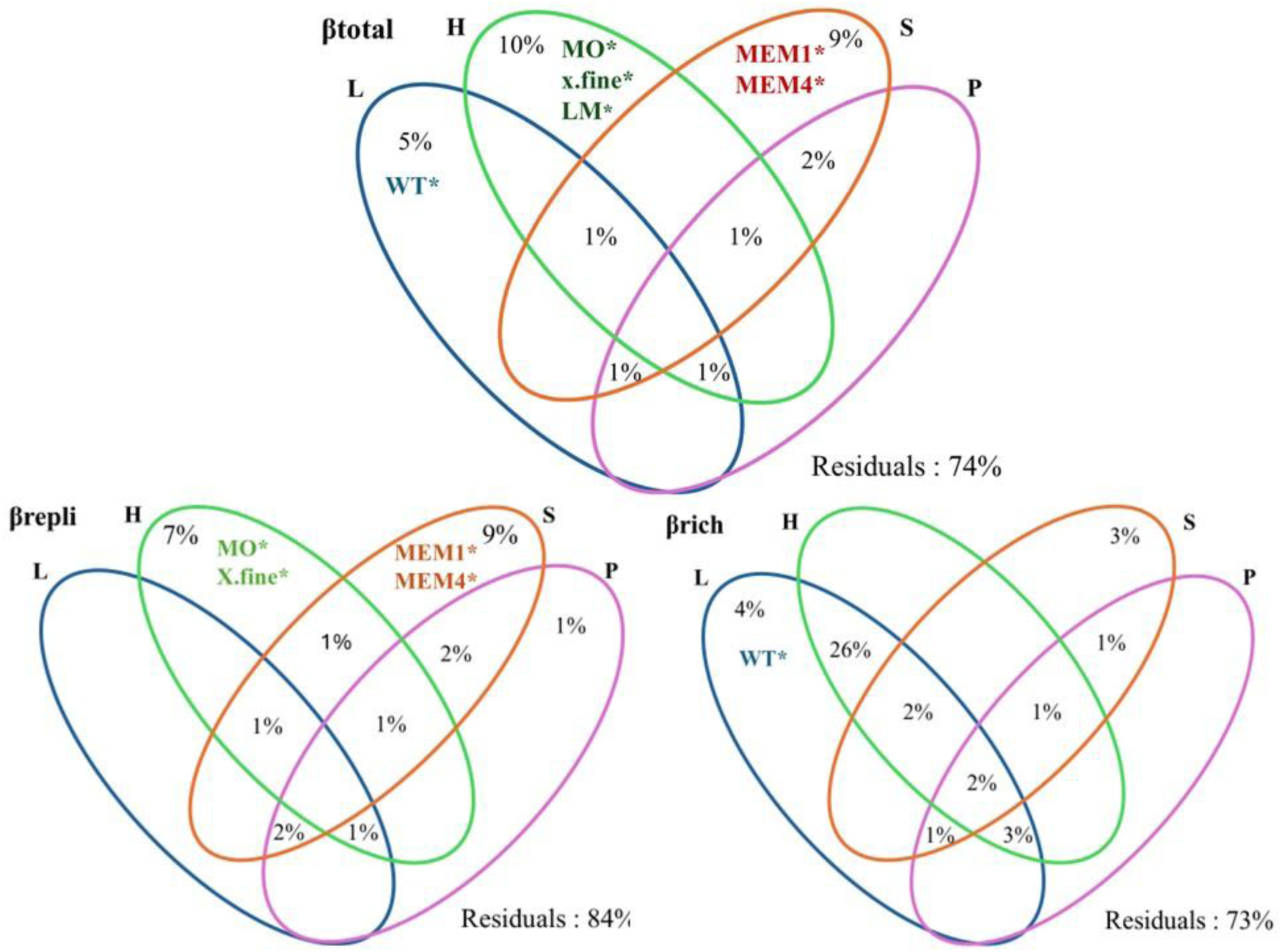
Venn diagrams representing the relative contribution of microhabitat (H), limnological (L), spatial (S), and landscape (P) variables in explaining taxonomic beta diversity: (a) total beta diversity (β_total), (b) replacement component (β_repl), and (c) richness difference (β_rich) of EPT communities.

### Drivers of functional β-diversity of EPT communities

Total functional beta diversity of EPT (F-β_total) was significantly explained only by local environmental variables. Habitat-related variables (H) accounted for 11% of the variation (F = 2.048; p = 0.025), with sand percentage (x.sand) and depth as the main predictors. Similarly, limnological variables, including water temperature (WT) and dissolved oxygen (DO), explained 11% of the variation (F = 2.057; p = 0.017). The spatial component did not show a significant effect on total functional beta diversity (F = 1.087; p = 0.370) (Figure 5).

**Figure 5.**
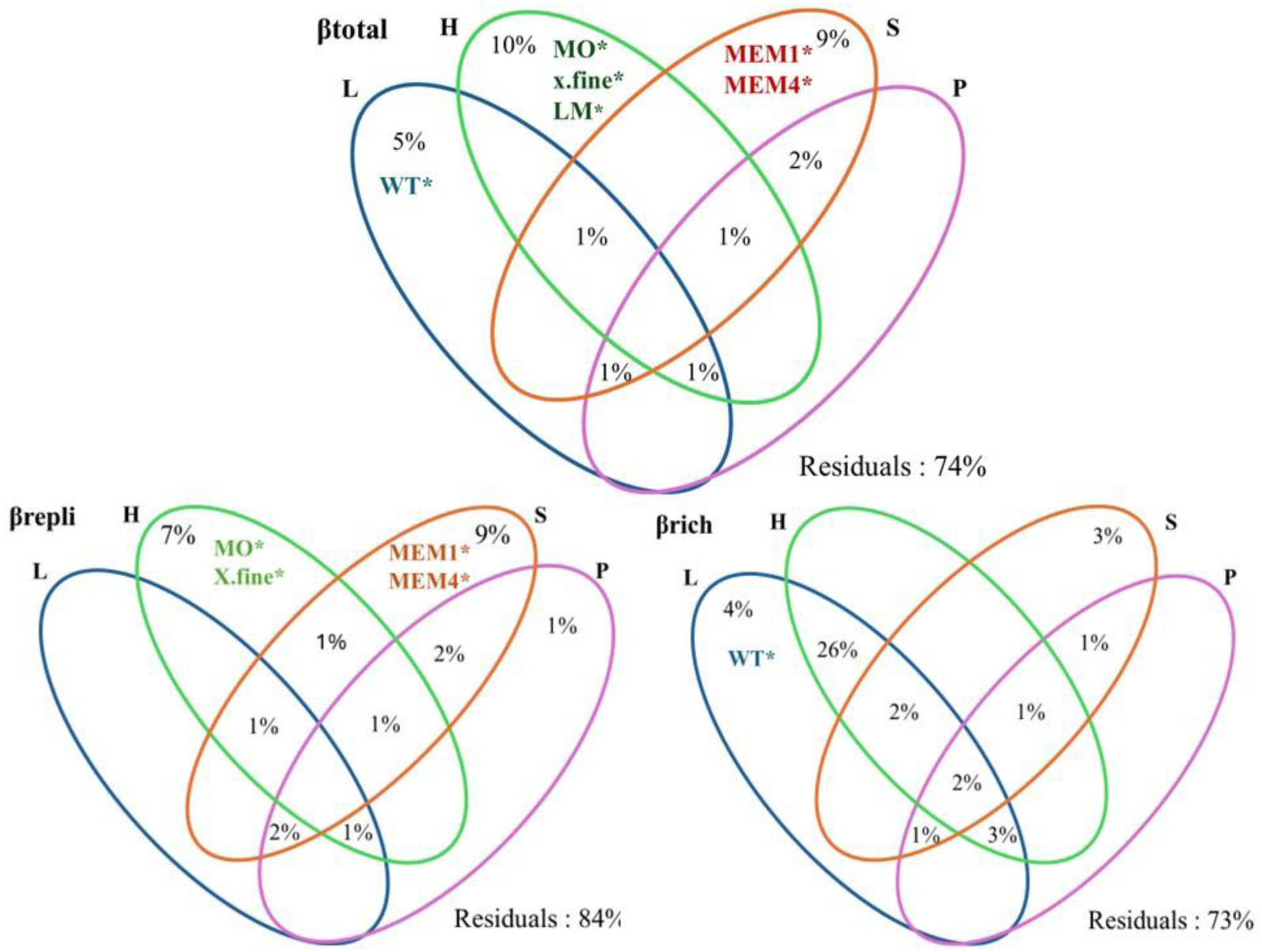
Venn diagrams representing the relative contribution of microhabitat (H), limnological (L), spatial (S), and landscape (P) variables in explaining functional beta diversity: (a) total functional beta diversity (F-β_total), (b) traits replacement (F-β_repl), and (c) functional richness difference (F-β_rich) of EPT communities.

None of the environmental, landscape, or spatial variables significantly explained the replacement- or richness-difference components of functional beta diversity. However, shared fractions showed substantial contributions, indicating that habitat, limnological, and landscape variables are spatially structured.

## Discussion

We observed that taxonomic beta diversity of EPT communities was predominantly structured by genus replacement, strongly associated with limnological variables, physical habitat characteristics, and spatial components. In contrast, functional beta diversity reflected the richness-difference component and was primarily explained by local environmental variables, supporting our hypotheses. This pattern reveals a decoupling between taxonomic and functional dimensions of biodiversity, indicating that distinct mechanisms govern genus composition and the distribution of functional traits in tropical lotic ecosystems. Similar patterns have been reported in Amazonian fish communities (e.g., Benone et al., 2020). Importantly, the high ecological integrity of the Panará Territory likely enhances the detectability of these patterns by reducing confounding anthropogenic disturbances, allowing underlying metacommunity processes to emerge more clearly.

The predominance of taxonomic replacement suggests that variation in EPT genus composition among streams in the Panará Territory occurs mainly along environmental and spatial gradients, rather than through genus gain or loss. This pattern is typical of relatively well-preserved environments, where environmental heterogeneity promotes speciation and species sorting processes, although still influenced by dispersal limitations (Leibold et al., 2004; Cunha & Juen, 2020; Brendonck et al., 2016). The significant influence of spatial components further supports the idea that drainage network configuration and the relative isolation of headwater streams contribute to structuring these communities (Nunes et al., 2020; Brown et al., 2010).

This effect may be particularly pronounced in headwater streams, which exhibit greater hydrological isolation and strong local environmental control over biotic assemblages (Brasil et al., 2016; Heino et al., 2015; Kulkarni et al., 2023). In these systems, organisms with passive aquatic dispersal depend heavily on habitat connectivity and immediate environmental conditions for dispersal (Barta et al., 2023; Heino et al., 2015). As a result, even small environmental differences among streams may lead to the replacement of ecologically specialized genera, reinforcing the observed taxonomic replacement. Thus, EPT community composition reflects, in a highly sensitive manner, the specific characteristics of each habitat, highlighting the interaction between spatial isolation and environmental filtering in shaping regional metacommunity structure (Heino et al., 2015; Barbosa et al., 2020).

In contrast, the predominance of functional richness-difference suggests the maintenance of a shared functional core across streams, despite taxonomic replacement. This result indicates high functional redundancy among EPT genera, such that taxonomic replacement occurs primarily among functionally equivalent taxa, which can be substituted with little or no impact on ecosystem processes (Nicácio et al., 2020; Hernández-Mendoza et al., 2024). This pattern is consistent with our expectation that functional beta diversity would be dominated by richness differences rather than replacement, reflecting the persistence of similar ecological strategies across sites. Importantly, this decoupling has direct implications for conservation, as taxonomic changes may not immediately translate into functional losses, although continued environmental change may progressively erode this redundancy over time.

The predominance of environmental filtering further promotes this compensatory dynamic, as functional traits tend to respond more consistently to environmental variation than taxonomic composition (Gothe et al., 2017). Global meta-analyses have shown that beta diversity components reflect distinct ecological processes: replacement is associated with environmental or historical gradients, whereas richness differences emerge when functional or taxonomic subsets are maintained under consistent environmental filters (Soininen et al., 2018). The predominance of functional richness differences observed here suggests that the relatively low environmental contrast among streams within a largely preserved landscape, combined with the structural conservation of these systems, selects for similar ecological strategies regardless of the taxonomic identity of the genera present (Castro et al., 2025). These results reinforce the idea that environmental filtering acts as a stabilizing mechanism on functional structure, even under conditions of high taxonomic turnover.

The selection of distinct predictors for beta diversity components reinforces this interpretation. Taxonomic replacement was associated with both spatial components and substrate characteristics, particularly organic matter and fine substrate fraction. The interaction between drainage network structure and benthic microhabitat heterogeneity drives genus replacement. These factors influence resource availability, substrate stability, and connectivity among streams, promoting distinct taxonomic assemblages across the landscape (Benone et al., 2017; Perez-Rocha et al., 2018; Branco et al., 2020). Although landscape variables showed limited direct effects, their spatial structuring suggests potential indirect influences mediated through local habitat conditions.

Conversely, functional richness was primarily explained by limnological and structural habitat variables, such as sand proportion, depth, and water temperature, which act as direct environmental filters on ecological and morphological traits. Thus, while taxonomic replacement reflects spatial reorganization of communities, functional patterns respond to environmental gradients that impose consistent constraints on viable strategies, reinforcing the role of species sorting in structuring these metacommunities (Heino & Tolonen, 2017; Gianuca et al., 2018; Hill et al., 2019).

This taxonomic–functional decoupling is consistent with the insurance hypothesis (Yachi & Loreau, 1999; Lansac-Tôha et al., 2022), as functional redundancy may buffer the effects of taxonomic replacement on community functional organization. In systems where species sorting consistently acts on functional traits, taxonomic identity may vary regionally while the functional spectrum remains relatively stable. Functional redundancy may therefore act as a buffering mechanism against taxonomic replacement, helping maintain ecosystem processes even under high species turnover. Contrasting patterns between taxonomic and functional diversity have also been observed in other tropical aquatic systems, where community structure results from the interaction between environmental filters and spatial processes, albeit with varying contributions of turnover and nestedness components (e.g., Hernández-Mendoza et al., 2024).

The importance of hydrological connectivity and spatial dynamics in structuring tropical aquatic communities has been widely documented. Variation in connectivity among streams can modulate dispersal and persistence of aquatic organisms, influencing taxonomic replacement, while consistent environmental filters maintain functional convergence (Lopez-Delgado et al., 2020). Even in contexts of low anthropogenic disturbance, subtle differences in connectivity can generate strong spatial patterns in community composition.

From a theoretical perspective, our results suggest that functional redundancy among EPT genera acts as a buffering mechanism against taxonomic replacement. However, this apparent stability does not imply invulnerability. Future changes in environmental filters or regional connectivity may simultaneously affect multiple functionally redundant genera, potentially leading to abrupt functional losses. Thus, the decoupling between taxonomic and functional dimensions highlights the complexity of community assembly processes in Amazonian streams and reinforces the need for multidimensional approaches to understand biodiversity dynamics and anticipate responses to environmental change. In this context, integrating taxonomic and functional perspectives is essential for improving biodiversity assessments and informing more effective conservation strategies. These findings also highlight the value of collaborative research approaches for generating robust ecological data in remote and underrepresented regions of the Amazon. Furthermore, the participatory approach adopted here demonstrates the potential of co-produced knowledge to enhance both scientific understanding and local engagement in biodiversity monitoring.

## Conclusion

The patterns observed in this study demonstrate that streams within Indigenous territories are key systems for understanding the processes that structure Amazonian aquatic biodiversity. The predominance of genus-level replacement in taxonomic beta diversity, coupled with differences in functional richness, indicates that these environments sustain high taxonomic heterogeneity without compromising the functional integrity of communities, reflecting the consistent action of environmental filters under conditions of high ecological integrity.

These findings reinforce the view that Indigenous territories function as reservoirs of biodiversity and functional stability, offering a unique opportunity to investigate ecological mechanisms under low direct anthropogenic pressure. Moreover, by revealing patterns of aquatic biodiversity organization in environments historically underrepresented in scientific research, this study highlights the urgency and importance of expanding investigations in freshwater systems within Indigenous territories, not only to advance ecological theory but also to recognize their fundamental role in preserving unique environments, maintaining essential ecosystem processes, and conserving Amazonian biodiversity across multiple scales. Protecting these systems is therefore critical for maintaining both biodiversity and the ecological processes that sustain resilience in Amazonian freshwater ecosystems.

## Acknowledgments

This study was financially supported by Conservation International (Grant/Project N° 115395 – Xingu-HP), which funded the research project and the scholarship granted to Jady Santos. We are deeply grateful for this support, which made fieldwork, data processing, and analyses possible. This study was also conducted within the framework of the Programa de Pesquisa em Biodiversidade da Amazônia Oriental (PPBio AmOr - process 441257/2023–2). We thank the Panará Indigenous Territory and its community members for authorizing and supporting this research, and for their collaboration throughout the study. We also acknowledge the relevant environmental authorities for granting the necessary research permits. JCMF thanks CAPES for the doctoral student scholarship (Process No. 88887.176346/2025-00). JSB and GMC are thankful to the National Council for Scientific and Technological Development (CNPq) for postdoctoral fellowships (processes 151038/2024-4 and 88887.939579/2024-00, respectively). JMRP is thankful to Hydro Barcarena for a postdoctoral fellowship associated with the project “PROJ. 4802 – Avaliação de Biota Aquática e Atributos Funcionais de Planta de Barcarena”. LFAM, LJ, RL and TSM is grateful for the research productivity grant (processes 302881/2022-1, 304710/2019-9, 312786/2025–5 and 311835/2023-6). This study was also supported by the Coordenação de Aperfeiçoamento de Pessoal de Nível Superior – Brazil (CAPES), Financing Code 001, through doctoral research fellowships awarded to LPO (Process No. 88887.939430/2024–00). We appreciate the support of the projects INCT Sínteses da Biodiversidade Amazônica (process 406767/2022–0), Advanced Research-Action Center for the Conservation and Recovery of the Amazon Ecosystem (CAPACREAM - process 444350/2024-1), and Programa de Pesquisa de Longa Duração (PELD-AmOr - process 445970/2024–3). We are especially grateful to the Biota Aquatic team from the Laboratório de Ecologia e Conservação and the Laboratório de Ecologia de Produtores Primários for their scientific and logistical support. We appreciate the support of the projects INCT Sínteses da Biodiversidade Amazônica process 406767/2022–0) and Programa de Pesquisa em Biodiversidade da Amazônia Oriental - PPBIO AmOr (process 441257/2023–2) and PELD-AmOr (process 445970/2024–3). We sincerely thank Panará Indigenous people and all collaborators who assisted during field sampling and laboratory procedures.

## Notes

### Competing Interest Statement

The authors have declared no competing interest.

## References

Azevedo-Santos, V. M., Frederico, R. G., Fagundes, C. K., Pompeu, P. S., Pelicice, F. M., Padial, A. A., Nogueira, M. G., Fearnside, P. M., Lima, L. B., Daga, V. S., Oliveira, F. J. M., Vitule, J. R. S., Callisto, M., Agostinho, A. A., Esteves, F. A., Lima-Junior, D. P., Magalhães, A. L. B., Sabino, J., Mormul, R. P., Grasel, D., Zuanon, J., Vilella, F. S., & Henry, R. (2019). Protected areas: A focus on Brazilian freshwater biodiversity. Diversity and Distributions 25:442–448.

Barbosa, D. D. A., Brasil, L. S., Azevêdo, C. A. S. D., & Lima, L. R. C. (2020). The role of spatial and environmental variables in shaping aquatic insect assemblages in two protected areas in the transition area between Cerrado and Amazônia. Biota Neotropica 20: e20190932.

Barrilli, G. H. C., do Vale, J. G., Chahad-Ehlers, S., Verani, J. R., & Branco, J. O. (2024). Spatial patterns of beta diversity in marine benthic assemblages from coastal areas of southern Brazil and their implications for conservation. Estuarine, Coastal and Shelf Science 297:108603.

Barta, B., Szabó, A., Szabó, B., Ptacnik, R., Vad, C. F., & Horváth, Z. (2023). How pondscapes function: Connectivity matters for biodiversity even across small spatial scales in aquatic metacommunities. Ecography 2024: e06960.

Bêche, L. A., & Statzner, B. (2009). Richness gradients of stream invertebrates across the USA: Taxonomy- and trait-based approaches. Biodiversity and Conservation 18:3909–3930.

Benone, N. L., Leal, C. G., dos Santos, L. L., Mendes, T. P., Heino, J., & de Assis Montag, L. F. (2020). Unraveling patterns of taxonomic and functional diversity of Amazon stream fish. Aquatic Sciences 82:75.

Blanchet, F. G., Legendre, P., & Borcard, D. (2008). Forward selection of explanatory variables. Ecology 89:2623–2632.

Branco, C. C., Bispo, P. C., Peres, C. K., Tonetto, A. F., Krupek, R. A., Barfield, M., & Holt, R. D. (2020). Partitioning multiple facets of beta diversity in a tropical stream macroalgal metacommunity. Journal of Biogeography 47:1765–1780.

Brasil, L. S., Dias-Silva, K., Jung, A., Oliveira, J. C. A., Sabino, U., & Vieira, T. B. (2016). Environment, space or connectivity: Which are the driving forces on aquatic insect assemblages in impounded streams? Entomotrópica 31:155–166.

Brasil, L. S., Luiza-Andrade, A., Calvão, L. B., Dias-Silva, K., Faria, A. P. J., Shimano, Y., Oliveira-Junior, J. M. B., Cardoso, M. N., & Juen, L. (2020). Aquatic insects and their environmental predictors: A scientometric study focused on environmental monitoring in lotic environments. Environmental Monitoring and Assessment 192:194.

Borcard, D., & Legendre, P. (2002). All-scale spatial analysis of ecological data by means of principal coordinates of neighbour matrices. Ecological Modelling, 153(1–2), 51–68.

Brown, B. L., & Swan, C. M. (2010). Dendritic network structure constrains metacommunity properties in riverine ecosystems. Journal of Animal Ecology 79:571–580.

Cadotte, M. W., Carscadden, K., & Mirotchnick, N. (2011). Beyond species: functional diversity and the maintenance of ecological processes and services. Journal of Applied Ecology, 48(5), 1079–1087.

Callisto, M. ., Solar, R. ., Rocha, A. S. da ., Paz, A. A. da ., Dolabela, B. M. ., Felisberto, B. ., Costa, E. C. da S. ., Eller, E. E. de O. ., Castro, H. F. L. de ., Gerheim, I. ., Lombello, J. C. ., Madureira, K. H. ., Souza, L. C. G. de ., Senna, N. ., Marques, R. ., Caffaro, R. M. ., Otuki, S. A. P. ., Santos, G. M. dos ., Amaral, P. H. M. do ., Carmo, F. F. do ., Kamino, L. H. Y. ., Linares, M. S. ., Ferraz, V. S. ., & Nunes, T. (2023). Avaliação ecológica rápida de qualidade de água e bioindicadores bentônicos no Parque Nacional da Serra do Gandarela, Minas Gerais. Revista Espinhaço.

Carvalho, R. L., Resende, A. F., Barlow, J., França, F. M., Moura, M. R., Maciel, R., Alves-Martins, F., Shutt, J., Nunes, C. A., Elias, F., Silveira, J. M., Stegmann, L., Baccaro, F. B., Juen, L., Schietti, J., Aragão, L., Berenguer, E., Castello, L., Costa, F. R. C., Guedes, M. L., Leal, C. G., Lees, A. C., Isaac, V., Nascimento, R. O., Phillips, O. L., Schmidt, F. A., ter Steege, H., Vaz-de-Mello, F., Venticinque, E. M., Vieira, I. C. G., Zuanon, J., The Synergize Consortium, & Ferreira, J. (2023). Pervasive gaps in Amazonian ecological research. Current Biology 33:3495–3504.e4.

Castro, D. M., do Amaral, P. H., van den Berg, E., Hughes, R. M., & Callisto, M. (2025). Spatial and temporal taxonomic and functional beta diversity of macroinvertebrate assemblages along a tropical dammed river. Aquatic Sciences 87:35.

Castro, D. M., da Silva, P. G., Solar, R., & Callisto, M. (2020). Unveiling patterns of taxonomic and functional diversities of stream insects across four spatial scales in the neotropical savanna. Ecological Indicators 118:106769.

Ceron, Y. M. M., & de Almeida Silva, T. T. (2023). Comunidades indígenas, meio ambiente e território: os casos dos territórios Raposa Serra do Sol no Brasil e do Parque Nacional Natural El Cocuy na Colômbia. Revista de Direito Ambiental e Socioambientalismo, 9(1), 66–85.

Chiaravalloti, R.M., Tomas, W.M., Leimgruber, P., Akre, T., & Giordano, A.J (2025). Science and Practice in the Pantanal: From crisis to hope. Conservation Science and Practice, 7(7).

Cunha, E. J., & Juen, L. (2020). Environmental drivers of the metacommunity structure of insects on the surface of tropical streams of the Amazon. Austral Ecology 45:586–595.

Dray, S., Blanchet, G., Borcard, D., Guenard, G., Jombart, T., Larocque, G., & Wagner, H. H. (2025). adespatial: Multivariate multiscale spatial analysis (R package version 0.3-28).

Dolédec, S., & Bonada, N. (2013). So what? Implications of biodiversity loss for ecosystem functioning. In S. Sabater, A. Elosegi, & R. Ludwig (Eds.), River conservation: challenges and opportunities (pp. 169–192). Fundación BBVA.

Domínguez, E. (2006). Ephemeroptera of South America (Vol. 2). Pensoft Publishers.

Façanha, B. L., Carvalho, R. L., Almeida, R. P., França, F. M., Sousa, J. R., Esposito, M. C., & Juen, L. (2026). Accessibility drives research efforts on Amazonian sarcosaprophagous flies. Proceedings of the Royal Society B: Biological Sciences 293:2064.

Faria, A. P. J., Ligeiro, R., Callisto, M., & Juen, L. (2017). Response of aquatic insect assemblages to the activities of traditional populations in eastern Amazonia. Hydrobiologia 802:39–51.

Faria, A. P. J., Ligeiro, R., Calvao, L. B., Giam, X., Leibold, M. A., & Juen, L. (2024). Land use types determine environmental heterogeneity and aquatic insect diversity in Amazonian streams. Hydrobiologia, 851(2), 281–298.

Faria, A. P. J., Paiva, C. K. S., Calvão, L. B., Cruz, G. M., & Juen, L. (2021). Response of aquatic insects to an environmental gradient in Amazonian streams. Environmental Monitoring and Assessment 193:763.

Faria, A. P. J., Ligeiro, R., Callisto, M., & Juen, L. (2017). Response of aquatic insect assemblages to the activities of traditional populations in eastern Amazonia. Hydrobiologia, 802(1), 39–51.

Gianuca, A. T., Engelen, J., Brans, K. I., Hanashiro, F. T. T., Vanhamel, M., van den Berg, E. M., Souffreau, C., & De Meester, L. (2018). Taxonomic, functional and phylogenetic metacommunity ecology of cladoceran zooplankton along urbanization gradients. Ecography 41:183–194.

Guerrero-Moreno, M. A., Silva, E. C., Oliveira, F. A., Nascimento, A. C. L., Michelan, T. S., Dias-Silva, K., Teodósio, M. A., Moura Jr, J. F., Oliveira-Junior, J. M. B., & Juen, L. (2025). Use of aquatic organisms as flagship species in selecting priority areas for conservation. Water Biology and Security:100509.

Grönroos, M., Heino, J., Siqueira, T., Landeiro, V. L., Kotanen, J., & Bini, L. M. (2013). Metacommunity structuring in stream networks: roles of dispersal mode, distance type, and regional environmental context. Ecology and Evolution, 3(13), 4473–4487.

Hamada, N. (2014). Insetos aquáticos na Amazônia brasileira: taxonomia, biologia e ecologia.

Heino, J., Melo, A. S., Siqueira, T., Soininen, J., Valanko, S., & Bini, L. M. (2015). Metacommunity organisation, spatial extent and dispersal in aquatic systems: Patterns, processes and prospects. Freshwater Biology 60:845–869.

Heino, J., Schmera, D., & Erős, T. (2013). A macroecological perspective of trait patterns in stream communities. Freshwater Biology, 58(8), 1539–1555.

Hepp, L. U., Milesi, S. V., Picolotto, R. C., Decian, V. S., Restello, R. M., Huiñocana, J. S., & Albertoni, E. F. (2023). Agriculture affects functional diversity of aquatic insects in Subtropical Atlantic Forest streams. Acta Limnologica Brasiliensia, 35, e31.

Hernández-Mendoza, L. C., Escalera-Vázquez, L. H., Vega-Cendejas, M. E., Núñez-Lara, E., Chiappa-Carrara, X., & Arceo-Carranza, D. (2024). Functional and taxonomic β diversity in fish assemblages is structured by turnover in a tropical coastal lagoon. Environmental Biology of Fishes 107:1219–1234.

Hill, M. J., Heino, J., White, J. C., Ryves, D. B., & Wood, P. J. (2019). Environmental factors are primary determinants of different facets of pond macroinvertebrate alpha and beta diversity in a human-modified landscape. Biological Conservation 237:348–357.

Intergovernmental Panel on Climate Change (IPCC) (2021). Climate change 2021: The physical science basis. Cambridge University Press, Cambridge.

Instituto Socioambiental, Associação Iakiô. 2025. Plano de gestão territorial e ambiental da Terra Indígena Panará = Kâprëpa Mï Hokïrähë Kypa Më, Aty Hä Kja Panäran Jö. Instituto Socioambiental, São Paulo.

Juen, L., Silva, F. S., Santos, F. M. B., Silva, B. L., Pérez, J. M. R., Junior, J. M. B. O., & Silva, K. D. (2025). Protocolo de coleta para inventário de insetos aquáticos na Amazônia no sistema RAPELD com ênfase em Ephemeroptera, Plecoptera, Trichoptera, Odonata e Heteroptera. EDUCAmazônia, 18, 18.

Juen, L., Cunha, E. J., Carvalho, F. G., Ferreira, M. C., Begot, T. O., Andrade, A. L., & Montag, L. F. A. (2016). Effects of oil palm plantations on the habitat structure and biota of streams in Eastern Amazon. River Research and Applications, 32(10), 2081–2094.

Junk, W. J., Bayley, P. B., & Sparks, R. E. (1989). The flood pulse concept in river-floodplain systems. Canadian Special Publication of Fisheries and Aquatic Sciences, 106, 110–127.

Laliberté, E., Legendre, P., & Shipley, B. (2014). FD: measuring functional diversity from multiple traits, and other tools for functional ecology. R package version 1.0–12.

Lansac-Tôha, F. M., Heino, J., Bini, L. M., Peláez, O., Baumgartner, M. T., Quirino, B. A., & Velho, L. F. M. (2022). Cross-taxon congruence of taxonomic and functional beta-diversity facets across spatial and temporal scales. Frontiers in Environmental Science, 10, 903074.

Laureto, L. M. O., Cianciaruso, M. V., & Samia, D. S. M. (2015). Functional diversity: an overview of its history and applicability. Natureza & Conservação, 13, 112–116.

Legendre, P., & Gallagher, E. D. (2001). Ecologically meaningful transformations for ordination of species data. Oecologia, 129(2), 271–280.

Legendre, P., Borcard, D., & Peres-Neto, P. R. (2005). Analyzing beta diversity: partitioning the spatial variation of community composition data. Ecological Monographs, 75(4), 435–450.

Leibold, M. A., Holyoak, M., Mouquet, N., Amarasekare, P., Chase, J. M., Hoopes, M. F., & Gonzalez, A. (2004). The metacommunity concept: a framework for multi-scale community ecology. Ecology Letters, 7(7), 601–613.

Lima, M., Firmino, V. C., de Paiva, C. K. S., Juen, L., & Brasil, L. S. (2022). Land use changes disrupt streams and affect the functional feeding groups of aquatic insects in the Amazon. Journal of Insect Conservation, 26(2), 137–148.

Malhi, Y., Roberts, J. T., Betts, R. A., Killeen, T. J., Li, W., & Nobre, C. A. (2008). Climate change, deforestation, and the fate of the Amazon. Science, 319(5860), 169–172.

Mammola, S., Carmona, C. P., Guillerme, T., & Cardoso, P. (2021). Concepts and applications in functional diversity. Functional Ecology, 35(9), 1869–1885.

Mateo-Vega, J., Potvin, C., Monteza, J., Bacorizo, J., Barrigón, J., Barrigón, R., & Meyer, C. (2017). Full and effective participation of indigenous peoples in forest monitoring for reducing emissions from deforestation and forest degradation (REDD+): trial in Panama’s Darién. Ecosphere, 8(2), e01635.

Maués-Silva, R., Barbosa Oliveira-Junior, J. M., Cruz, G. M. D., & Brasil, L. S. (2024). Evaluation of the environmental quality monitoring protocol for Amazonian streams: a systematic review. Revista Ambiente & Água, 19, e2967.

Mouchet, M. A., Villéger, S., Mason, N. W., & Mouillot, D. (2010). Functional diversity measures: an overview of their redundancy and their ability to discriminate community assembly rules. Functional Ecology, 24(4), 867–876.

Monteles, J. S., Gerhard, P., Ferreira, A., & Sonoda, K. C. (2021). Agriculture impacts benthic insects on multiple scales in the Eastern Amazon. Biological Conservation 255:108998.

Moutinho, Z., Rocha, D. B., & Bechelany, F. (2025). A vulnerabilidade do povo indígena Panará à contaminação das águas superficiais. AMBIENTES: Revista de Geografia e Ecologia Política, 7(2).

Mozzaquattro, L. B., Dala-Corte, R. B., Becker, F. G., & Melo, A. S. (2020). Effects of spatial distance, physical barriers, and habitat on a stream fish metacommunity. Hydrobiologia 847:3039–3054.

Nessimian, J. L., Venticinque, E., Zuanon, J., De Marco, P., Gordo, M., Fidelis, L., Batista, J. D., & Juen, L. (2008). Land use, habitat integrity, and aquatic insect assemblages in Central Amazonian streams. Hydrobiologia 614:117–131.

Nicacio, G., Cunha, E. J., Hamada, N., & Juen, L. (2020). Contrasting beta diversity and functional composition of aquatic insect communities across local to regional scales in Amazonian streams. *bioRxiv*:2020.09.14.297077.

Nunes, C. A., Castro, F. S., Brant, H. S., Powell, S., Solar, R., Fernandes, G. W., & Neves, F. S. (2020). High temporal beta diversity in an ant metacommunity, with increasing temporal functional replacement along the elevational gradient. Frontiers in Ecology and Evolution 8:571439.

Oksanen, J., Blanchet, F. G., Kindt, R., Legendre, P., Minchin, P. R., O’hara, R. B., & Wagner, H. (2019). vegan: Community ecology package (version 2.5-6). Available from https://CRAN.R-project.org/package=vegan

Osborne, T., Cifuentes, S., Dev, L., Howard, S., Marchi, E., Withey, L., & Silva, M. S. R. (2024). Climate justice, forests, and Indigenous peoples: Toward an alternative to REDD+ for the Amazon. Climatic Change 177:128.

Peck, D. V., Herlihy, A. T., Hill, B. H., Hughes, R. M., Kaufmann, P. R., Klemm, D. J., & Cappaert, M. R. (2006). Environmental monitoring and assessment program–surface waters western pilot study: field operations manual for wadeable streams. U.S. Environmental Protection Agency.

Pereira, F. F., Pellin, Â., Dias, L., Silva, M., Lehmann, D., Bernardes, V., … & Tófoli, C. (2022). Percepção do Conselho acerca do Monitoramento Participativo da Biodiversidade para a Gestão das Unidades de Conservação da Amazônia. Biodiversidade Brasileira, 12(5), 151–166.

Peres-Neto, P. R., Legendre, P., Dray, S., & Borcard, D. (2006). Variation partitioning of species data matrices: Estimation and comparison of fractions. Ecology 87:2614–2625.

Perez-Rocha, M., Bini, L. M., Domisch, S., Tolonen, K. T., Jyrkänkallio-Mikkola, J., Soininen, J., Hjort, J., & Heino, J. (2018). Local environment and space drive multiple facets of stream macroinvertebrate beta diversity. Journal of Biogeography 45:2744–2754.

Pignati, W. A., Lima, F. A. N. D. S., Lara, S. S. D., Correa, M. L. M., Barbosa, J. R., Leão, L. H. D. C., & Pignatti, M. G. (2017). Distribuição espacial do uso de agrotóxicos no Brasil: uma ferramenta para a Vigilância em Saúde. Ciência & Saúde Coletiva, 22, 3281–3293.

Podani, J., & Schmera, D. (2011). A new conceptual and methodological framework for exploring and explaining pattern in presence–absence data. Oikos 120:1625–1638.

Poff, N. L., Olden, J. D., Vieira, N. K., Finn, D. S., Simmons, M. P., & Kondratieff, B. C. (2006). Functional trait niches of North American lotic insects: traits-based ecological applications in light of phylogenetic relationships. Journal of the North American Benthological Society, 25(4), 730–755.

Santos, N. B. B., Cruz, G. M., Monteles, J. S., Faria, A. P. J., Firmino, V. C., Shimano, Y., Ferreira, V. R. S., Luiza-Andrade, A., Salles, F. F., Castro, D. M. P., Quinteiro, F. B., Lima, L. R. C., Dias, L. G., Pes, A. M. O., Hamada, N., & Juen, L. (2024). Database of immature stage traits of Ephemeroptera, Plecoptera, and Trichoptera (EPT) genera for the Amazon. Aquatic Sciences 86:35.

Shimano, Y., Cardoso, M., & Juen, L. (2018). Estudos ecológicos de efemerópteros (Insecta, Ephemeroptera): O esforço amostral pode ser reduzido sem perder informações taxonômicas e ecológicas essenciais? Acta Amazonica 48:137–145.

Silva Santos, J. V. A., Lima, M., Monteles, J. S., Carrera, D. L. R., de Faria, A. P. J., Brasil, L. S., & Juen, L. (2023). Assessing physical habitat structure and biological condition in eastern Amazonia stream sites. Water Biology and Security 2:100132.

Silva, E. C., Guerrero-Moreno, M. A., Oliveira, F. A., Dias-Silva, K., Juen, L., Moura Junior, J. F., Carvalho, F. G., & Oliveira-Junior, J. M. B. (2025). Socio-environmental conflicts and traditional communities in protected areas: A scientometric analysis. Journal for Nature Conservation 86:126936.

Soininen, J., Heino, J., & Wang, J. (2018). A meta-analysis of nestedness and turnover components of beta diversity across organisms and ecosystems. Global Ecology and Biogeography 27:96–109.

Soininen, J., Lennon, J. J., & Hillebrand, H. (2007). A multivariate analysis of beta diversity across organisms and environments. Ecology, 88(11), 2830–2838.

Southwood, T. R. (1977). Habitat, the templet for ecological strategies? Journal of Animal Ecology 46:337–365.

Rocha, D. F. B. (2024). Quando o réu é o Estado: revisitando a ação indenizatória dos Panará à luz do direito à memória, verdade, reparação e responsabilização para não-repetição.

Rosenberg, D. M., & Resh, V. H. (1993). Freshwater biomonitoring and benthic macroinvertebrates. Chapman & Hall.

Thorp, J. H., & Covich, A. P. (2010). Ecology and classification of North American freshwater invertebrates (3rd ed.). Academic Press, Amsterdam.

Tockner, K., Pusch, M., Borchardt, D., & Lorang, M. S. (2010). Multiple stressors in coupled river–floodplain ecosystems. Freshwater Biology 55:135–151.

Villéger, S., Grenouillet, G., & Brosse, S. (2013). Decomposing functional β-diversity reveals that low functional β-diversity is driven by low functional turnover in European fish assemblages. Global Ecology and Biogeography, 22(6), 671–681.

Ward, J. V. (1989). The four-dimensional nature of lotic ecosystems. Journal of the North American Benthological Society 8:2–8.

Yachi, S., & Loreau, M. (1999). Biodiversity and ecosystem productivity in a fluctuating environment: The insurance hypothesis. Proceedings of the National Academy of Sciences 96:1463–1468.

